# Genome-wide exploration of metabolic-based pyrethroid resistance mechanism in *Helicoverpa armigera*

**DOI:** 10.1101/2023.12.18.572109

**Authors:** Juil Kim, Md-Mafizur Rahman, Changhee Han, Jungwon Jeon, Min Kwon, Si Hyeock Lee, Celso Omoto

## Abstract

To elucidate the deltamethrin resistance mechanism in *Helicoverpa armigera*, we explored mutations at the deltamethrin target site, genomic level variations between insecticide-susceptible and -resistant strains, and differences in gene expression patterns between the strains. Known pyrethroid resistance-associated point mutations within the voltage-gated sodium channel were undetected in the cDNA and gDNA of resistant strains or field populations. The whole-genome *de novo* assembly of a Korean resistant strain was performed (GCA_026262555.1), and 13 genomes of susceptible and resistant individuals were re-sequenced using field populations. Approximately 3,369,837 variants (SNPs and indels) were compared with our reference *H. armigera* genome, and 1,032,689 variants were identified from open reading frames. A resistance-specific CYP3 subfamily gene with five variants (CYP321A1v1–v5) was identified in the resistant strains, indicating the potential role of these variants in resistance. RNA-seq analysis identified 36,720 transcripts from 45 Illumina RNA-seq datasets of the fat body, gut, and the rest of the body. Differential gene expression analysis revealed some differently overexpressed detoxification enzyme genes in the resistant strains, particularly cytochrome P450 genes. This finding was consistent with the results of bioassay tests using PBO-based synergists, further supporting the role of detoxification enzymes in resistance. Therefore, *H. armigera* may acquire deltamethrin resistance through a combination of actions, including the overexpression of various detoxification enzymes, such as CYP3 subfamilies (CYP321A5) and cuticular proteins. The five variants of CYP321A subfamily identified in this study may serve as a basis for understanding insecticide resistance at the molecular level and can be applied as diagnostic markers for resistance.

**Key Messages:**

- Known resistance-related mutations were undetected in all the resistant strains or field populations.
- No specific resistance-associated variations were identified at the genomic level.
- The expression pattern of the CYP3 subfamily genes was strongly correlated with the level of resistance.
- Genes other than CYP337B3 are also involved in the development of high-level resistance.
- Resistance developed as a result of changes in the expression of detoxification genes rather than target site modification through mutation.

## INTRODUCTION

The cotton bollworm [*Helicoverpa armigera* (Hübner)] is a destructive pest that affects a wide range of crops (Riaz et al. 2021). Widespread insecticide use in farming has led to the development of insecticide resistance in many insect pest species, including *H. armigera*. This phenomenon is considered among the most prominent examples of microevolution (Siddiqui et al. 2023). Intensive insecticide use against *H. armigera* has led to the development of resistance to most insecticides used for its control, with 891 cases of insecticide resistance to 55 active ingredients recorded for this species (Database 2023; Stavrakaki et al. 2023). Multiple factors contribute to the emergence of resistance, including biological, genetic, and operational factors (Sakine 2012). Among these, genetic factors are particularly important, as they play a crucial role in shaping resistance mechanisms in pest populations (Sakine 2012; Siddiqui et al. 2023). Identification of specific mutations responsible for insect resistance to certain insecticides, including pyrethroids, is essential (Liu et al. 2000; Shi et al. 2021). Two notable mutations, *kdr* (knockdown resistance) and *super-kdr*, occur in the voltage-gated sodium channel (VGSC) of insects, e.g., at positions 918 and 1,014 in the housefly (*Musca domestica*)(Davies et al. 2007). Previous studies have indicated that enhanced activity of detoxification enzymes, such as cytochrome P450 monooxygenase (CYP), carboxyl/choline esterase (CCE), and glutathione S-transferase (GST) (Chen et al. 2007; Joußen et al. 2012; Bai et al. 2019) and point mutations in the target channels of VGSC (Head et al. 1998; Hopkins and Pietrantonio 2010; Schmidt et al. 2010) may be associated with pyrethroid resistance in *H. armigera*. In *Helicoverpa zea*, three point mutations (V421A/G, I951V, and L1029H) are involved in pyrethroid resistance mechanisms (Hopkins and Pietrantonio 2010). Two additional mutations (D1556 and E1560) have been identified (Head et al. 1998) and functionally confirmed (Chen et al. 2017).

Single nucleotide polymorphisms (SNPs) and indels (insertions and deletions) appear to be associated with resistance mechanisms (Han et al. 2023). A high-quality chromosome-level whole insect genome could offer valuable insights into resistance mechanisms (Zhang et al. 2020). Various lepidopteran genome studies (Han et al. 2023; Kim et al. 2021), particularly those of *H. armigera,* have provided useful information to the scientific community, as demonstrated by earlier research (Zhang et al. 2010; Muller et al. 2021).

To gain insights into the molecular mechanisms underlying insecticide resistance, researchers have employed RNA deep sequencing technology (RNA-seq) (Trapnell et al. 2013). Lin et al. (2013) identified 1,215 genes that may be involved in chlorantraniliprole resistance in three field-resistant *Plutella xylostella* strains. Several of these genes were associated with calcium signaling, vascular smooth muscle contraction, and cardiac muscle contraction pathways, as well as the metabolism of xenobiotics including insecticides (Lin et al. 2013). Previous studies have suggested that P450 enzymes, encoded by a superfamily of genes, play crucial roles in the metabolism of both endogenous and exogenous substances in various organisms (Nauen et al. 2022). They contribute to the adaptability of pests to a wide range of insecticides. In *H. armigera*, multiple CYP genes have been identified as overexpressed and associated with deltamethrin insecticide resistance (Pittendrigh et al. 1997; Yang et al. 2006). Examples include CYP6B7 (Zhang et al. 2010), CYP9A12, CYP9A14, and CYP337B3 (Joußen et al. 2012; Rasool et al. 2014). However, the expression levels of these specific genes can vary among different *H. armigera* strains, and the contribution of a particular CYP gene to resistance can also differ. In addition, insect genomes encode several coding and non-coding RNAs that interact with and regulate gene expression, including that of CYPs, thereby influencing insect phenotypes (Yin et al. 2016; Satyavathi et al. 2017; Zhu et al. 2017; Cipolla et al. 2018).

An earlier study reported moderate-to-extremely high levels of resistance (over 20,000-fold) against pyrethroid insecticides in *H. armigera* populations (Durigan et al. 2017). Previous reports from Australia (Gunning et al. 1995), Spain, Turkey, Pakistan (Joußen et al. 2012), India (Kranthi et al. 1997), China (Ni et al. 2023), and Greece (Stavrakaki et al. 2023) have also documented moderate-to-high resistance levels against pyrethroids, demonstrating the ubiquity of pyrethroid resistance. The development of pyrethroid resistance in *H. armigera* has been associated with various mechanisms, including metabolic resistance, target-site resistance mutations, and reduced penetration (Gunning et al. 1995), among others. *H. armigera* is among the most commonly reported species in cases of insecticide resistance worldwide, demonstrating an evolved upregulation of metabolic resistance genes against various types of insecticides, including pyrethroids (Joußen et al. 2012; Rasool et al. 2014; Durigan et al. 2017). Metabolic resistance involves the regulation of gene expression by specific detoxification enzymes, including P450 (Pittendrigh et al. 1997; Durigan et al. 2017), CCE (Teese et al. 2010; Joußen et al. 2012), and cuticular proteins (CP) (Balabanidou et al. 2018).

Therefore, this study explored three potential mechanisms of deltamethrin resistance in *H. armigera*: (1) mutation of the target sites, (2) genomic variation, and (3) gene expression patterns. Multiple levels of investigation were employed to understand the underlying factors contributing to resistance. We focused on several key aspects to gain a comprehensive understanding of the resistance phenomenon in *H. armigera* (Cunningham and Zalucki 2014). Firstly, we performed *de novo* assembly of the whole genome of *H. armigera* using a draft genome as a reference. In addition, we analyzed 13 re-sequenced genome sets, including susceptible and resistant individuals with varying levels of resistance (low, medium, and high). Secondly, we used genome-wide pool-seq data alignment of VGSC genes to screen for target-site mutations. We examined the presence and frequency of point mutations in this population. Furthermore, genetic analysis focused on the five previously identified point mutations specifically associated with pyrethroid resistance, which are found within partial genes of VGSC (Head et al. 1998; Hopkins and Pietrantonio 2010; Chen et al. 2017). And finally, we utilized RNA-seq data from the susceptible and resistant strains (three categories: low, medium, and high resistance insects and tissues from each strain: fat, gut, and the rest of the body) to investigate the relationship between insecticide resistance and transcript expression.

## MATERIALS AND METHODS

### Insects

One susceptible (Australian susceptible, TWB-S) and three pyrethroid insecticide-resistant *H. armigera* strains (Australian resistant, TWB-R; Korean resistant, Kor-R; Brazilian resistant, bA43) were used in this study. The two TWB strains, originating in Toowoomba, Queensland, Australia, in 2003, have been maintained in the laboratory. The susceptible strain, TWB-S, lacks the CYP337B3 gene and was kept without insecticide exposure in the laboratory (Joußen et al. 2012), donated by an international collaborative research program (Project No. PJ011777). The resistance levels were categorized as follows: A) The low-resistant (TWB-R) strain is an isogenic strain of TWB-S, requiring an approximately 40-fold higher lethal dose (LD_50_) of fenvalerate (Joußen et al. 2012). This strain harbors the CYP337B3 gene, which is segregated from an identical population. B) The resistance of the Kor-R strain (collected from a cornfield in Pyeongchang, South Korea, 2013) to deltamethrin was higher than that of TWB-R (Kim et al. 2018). C) The highly pyrethroid-resistant strain bA43 (approximately 20,000-fold) was collected from soybean fields in Lous Eduardo Magalhaes, Brazil (Durigan et al. 2017). Because the bA43 strain showed an extremely high level of resistance to pyrethroids such as deltamethrin, it was difficult to determine the 100% mortality dose or concentration. Therefore, the LD_50_ or median lethal concentration (LC_50_) values could not be calculated. All susceptible and resistant strains (categorized as A, B, and C) were reared on a Bio-Serv artificial diet (F9772; Bio-Serv, Flemington, NJ, USA) at 26°C and 55% relative humidity with a 16:8 (L: D) photoperiod under previously reported conditions (Joußen et al. 2012).

### Bioassay

Technical grade deltamethrin (CAS Number: 52918-63-5, purity: 98.6%; PESTANAL, Sigma-Aldrich, St. Louis MO, USA) was used for the bioassay. Commercial products (emulsifiable concentrate, deltamethrin 1%; Bayer Crop Science, Monheim, Germany) were also used for selection and partially used for the synergy test. The bioassay assessed deltamethrin against the 3^rd^ instar larvae of *H. armigera,* comparing the insecticide-susceptible strain (TWB-S), insecticide-resistant strain (TWB-R), Korean resistant strain (Kor-R), and the Kor-R and TWB-R hybrid (male Kor-R × female TWB-R). Different deltamethrin doses were dissolved in acetone and topically applied to the dorsal part of the third-instar larvae (more than 30 individuals per dose) using a Hamilton syringe fixed in a Hamilton PB600-1 repeating dispenser (Hamilton, Reno, NV, USA). The larvae were maintained on a Bio-Serv diet under normal rearing conditions, as mentioned above. After 48 h, larval mortality rates were recorded.

Synergy tests were performed using 3^rd^ instar *H. armigera* larvae by administering deltamethrin at two doses (approximately LD_50_ and LD_10_) with or without the P450 synergist piperonyl butoxide (PBO; CAS Number: 51-03-6, Sigma-Aldrich) according to a previously reported method, with some modifications applied to *Spodoptera exigua* (Kim et al. 2021).

The probit model in SAS (SAS Institute 9.1, Cary, NC, USA) was used to estimate the concentration-based mortality after 2 days of insecticide treatment to determine the LC_50_ and 95% confidence limits. The susceptible species served as the standard reference for all comparisons of the tested insecticides, and the resistance ratios (RRs) and their associated 95% confidence intervals (CIs) were determined. The RR was computed by dividing the LC_50_ value of the population by that of the susceptible strain. Detailed sample information is provided in Table 1.

**Table 1.** Detailed sample information of insecticide-susceptible and -resistant strains and Korean field populations.

| No. | Sample code | Sample sources | Collection<br>year | Note | References |
| --- | --- | --- | --- | --- | --- |
| 1 | TWB3 | Susceptible strain from<br>Australia | 2003 | Susceptible strain | (Joußen et al. 2012) |
| 2 | bA33 | Resistant strain from<br>Brazil | 2013 | Resistant strain | (Durigan et al. 2017) |
| 3 | bA43 | Resistant strain from<br>Brazil | 2014 | Resistant strain | (Durigan et al. 2017) |
| 4 | Kor-R | Resistant strain from<br>Korea | 2015 | Resistant strain | (Kim et al. 2018) |
| 5 | BA-21 | Buan, Korea | 2021 | Field population | This study |
| 6 | DG-21 | Daegwallyeong, Korea | 2021 | Field population | This study |
| 7 | HN-21 | Haenam, Korea | 2021 | Field population | This study |
| 8 | HS-21 | Hoengseong, Korea | 2021 | Field population | This study |
| 9 | IJ-21 | Injae, Korea | 2021 | Field population | This study |
| 10 | JD-21 | Jindo, Korea | 2021 | Field population | This study |
| 11 | JE-21 | Jeongeup, Korea | 2021 | Field population | This study |
| 12 | PH-21 | Pohang, Korea | 2021 | Field population | This study |
| 13 | SW-20 | Suwon, Korea | 2020 | Field population | This study |

### VGSC mutation and CYP321A variant surveys

For the mutation sequence analysis, genomic DNA (gDNA) was individually extracted from each strain collected from the three levels of resistant populations and ten different field population locations across all populations using DNAzol (Molecular Research Center, Cincinnati, OH, USA), as previously described (Kim et al. 2021). Total RNA was extracted from the same strains and populations using a RNeasy Mini Kit (Qiagen, Hilden, Germany), following the manufacturer’s instructions. A TOPscript cDNA synthesis kit (Enzynomics, Daejeon, Korea) was used to generate cDNA. Both cDNA and gDNA were quantified using a NanoDrop spectrophotometer (NanoDrop Technologies, Wilmington, DE, USA).

Five previously reported non-synonymous mutations (V421A/G, I951V, L1029H, D1561V, and E1565C) were screened in the VGSC sequence. VGSC fragments were partially amplified on an Applied Biosystems ProFlex PCR system (Thermo Fisher Scientific, Waltham, MA, USA) using KOD FX polymerase (Toyobo Life Science, Osaka, Japan) with newly designed primer combinations (Table S1) and PCR conditions. The thermal conditions used for the touchdown PCR were as follows: initial denaturation at 94°C for 1 min; followed by 40 cycles of denaturation at 94°C for 20 s, annealing at 57–62°C for 20 s, extension at 68°C for 20 s; and a final extension at 68°C for 2 min followed by holding at 8°C until removal. Direct sequencing of PCR products was performed by Macrogen (Seoul, Korea). CYP321A variants were verified by sequencing using CYP321A universal primers and nested primer sets (Table S1) under the conditions described above. The loop-mediated isothermal amplification (LAMP) assay protocol was performed based on a previously reported method (Kim et al., 2021).

### Genome assembly and re-sequencing

For genome sequencing, gDNA was extracted from individual 5^th^ instar larvae of the Kor-R strain using a Quick DNA universal kit (Zymo Research, Irvine, CA, USA). The gDNA was validated and quantified using an Agilent 2200 TapeStation (Agilent Technologies, Santa Clara, CA, USA). The 29.1 Gb nanopore data (76.6×) underwent self-correction using the Canu assembler (version 1.71) and was then *de novo* assembled using SMARTdenovo (https://github.com/ruanjue/smartdenovo) with default parameters. The assembled contig sequences were polished once using Pilon (version 1.23) (Walker et al. 2014) with trimmed 15.8 Gb Illumina sequencing data (41.7×). The polished contig sequences were deposited in GenBank (GCA_017165865.1). The contig sequences were further scaffolded to chromosome-level assembled sequences using 48.7 Gb Hi-C data. Chromosome-level assembled sequences were deposited in GenBank (GCA_026262555.1). Repeat sequences were identified using RepeatModeler (version 1.0.11, http://www.repeatmasker.org/RepeatModeler.html) and RepeatMasker (version 4.0.9, http://www.repeatmasker.org/). Protein-coding genes were predicted in the genome sequence using an evidence-based annotation pipeline comprising MAKER3 (version 3.01.03) and EvidenceModeler (version 1.1.1). For transcriptome evidence, the RNA-seq data of *H. armigera* were used. Protein sequences from the reference genome Harm1.0 (GCF_002156985.1) served as protein evidence. Genome sequencing, assembly, and annotation were performed by Phyzen Co., Ltd. (Phyzen, Seongnam, Korea). Pairwise synteny between two genome datasets (*H. armigera* and *H. zea*) was investigated using LAST v1080 (LAST, RRID:SCR_006119) (Kiełbasa et al. 2011), and the Python-based package McScan (MCScan, RRID:SCR_017650) (Tang et al. 2008) in JCVI (Tang et al. 2015) was used to filter out tandem duplications and weak hits (Hill et al. 2019). Finally, the high-quality syntenic blocks were visualized using Circos v0.69-9 (Circos, RRID:SCR_011798) (Krzywinski et al. 2009).

For re-sequencing, gDNA was individually extracted from the larval stages of insecticide-susceptible and - resistant strains and field populations of *H. armigera* using a Quick DNA universal kit (Zymo Research). The quality and quantity of gDNA from 13 samples were examined using an Agilent 2200 TapeStation (Agilent Technologies). Paired-end (PE) sequencing libraries were constructed from the gDNA using a TruSeq DNA PCR free kit (Illumina) and subsequently sequenced on an Illumina HiSeqX platform (Illumina). Low-quality and duplicated reads, as well as adapter sequences, were removed using Trimmomatic (version 0.39) (Bolger et al. 2014) with default parameters. The trimmed high-quality reads were then mapped to the reference genome (GCA_026262555.1) using Burrows–Wheeler Aligner software (version 0.7.17) (Li and Durbin, 2009) with mem parameters. SAMtools software (version 1.11) (Li and Durbin 2009) was used to remove unmapped and secondary aligned reads and convert the mapping results into BAM format. Variation calling was performed using GATK (version 4.2) (McKenna et al. 2010), and then VCF files were generated. The identified SNPs and indels were annotated using SnpEff software (version 5.2e) (Cingolani et al. 2012).

### RNA-seq and differentially expressed gene (DEG) analyses

For RNA extraction, five larvae (5^th^ instar larvae within 12 h after molting) were collected in tubes, and each set of five larvae was considered one sample. Each insect was dissected into the fat body, gut, and the rest of the body. Additionally, the Kor-R strain was treated with a sublethal dose of deltamethrin (0.002 µg/larva) (designated as “Kor-T”) 24 h before dissection. According to the manufacturer’s protocol, 45 RNA-seq data samples were extracted from three tissue samples (three biological replicates of each insect), collected from three resistant strains (TWB-R, Kor-R/Kor-T, and bA43) and one susceptible strain (TWB-S) using a RNeasy Mini Kit (Qiagen). The integrity of the total RNA was validated and quantified using the same procedure (Agilent Technologies). Libraries were prepared using the TruSeq RNA Sample Prep Kit v2 (Illumina) and sequenced on the Hiseq4000 platform with the TruSeq 3000/4000 SBS Kit v3 (Macrogen, Seoul, Korea).

DEG analysis was performed in two ways: mapping to the reference genome (GCF_002156985.1, Harm_1.0) and mapping by merging all analyzed RNA-seq results to create the UniGene set. RNA-seq data were trimmed using Trimmomatic v0.36 (Bolger et al. 2014). Trimmed high-quality RNA-seq reads were mapped onto the reference gene dataset (GCA_026262555.1), and the transcripts per million and fragments per kilobase of transcript per million (FPKM) counts were calculated based on the RNA-seq data of the gene sequences using the RNA-seq by Expectation Maximization program ver. 1.2.9 (Robinson et al. 2010) with default parameters. A *p-*valueL<L0.05 and |log2 (fold change) | >2 were considered significantly differentially expressed transcripts. Statistical analysis was performed based on FPKM values. To compare expression patterns by strains or tissues, an MA plot (Robinson et al. 2010) was generated using the heatmap.2: function in the R package “gplots” (v3.1.1).

For a more integrated DEG analysis among all tested strains, because some strain-specific genes may be omitted from the reference genome database, the UniGene set was constructed using Trinity (ver. 2.12.0), CD-HIT-EST (ver. 4.8.1), and Transdecoder (ver. 5.5), based on the whole RNA-seq sequencing results. The DEG analysis was performed in the same manner as the expression analysis based on the reference genome mentioned above.

## RESULTS

### Bioassays and synergy tests to determine the causes of resistance

The LD_50_ of deltamethrin against the susceptible strain (TBW-S) was 0.0022 µg/larva (95% CI interval, 0.00122–0.00372 µg/larva), whereas that for the resistant strain (TBW-R) was 0.05818 µg/larva (95% CI, 0.04286–0.07430 µg/larva). The calculated RR^s^ was 26, indicating that the resistant strain was 26 times less susceptible to deltamethrin than the susceptible strain. The LD_50_ of deltamethrin against the Kor-R strain was 14,868-fold higher than that of the susceptible strain (TBW-S), indicating a remarkably elevated level of resistance. The RR^r^ values of the F1 hybrid and Kor-R were 265-(15.4443/0.0582) and 562-fold (32.7093/0.0582), respectively (Table 2). Notably, strains TBW-S and TBW-R differed in the presence or absence of CYP337B3, indicating its potential role in resistance. Although CYP337B3 may account for the differences between TBW-S and TBW-R, it is hypothesized that the high-level resistance observed in Kor-R cannot be solely attributed to CYP337B3, suggesting the involvement of additional factors. In addition, the F1 hybrid was produced by crossing a strain TWB-R female and a strain Kor-R male, considering maternal inheritance. This approach assumes that the resistance-related gene(s) in Kor-R are located on autosomes rather than on sex chromosomes. The observed resistance in the F1 hybrid suggests the potential transmission of metabolic genes that may affect the resistance of subsequent generations. The χ2, value suggests that a broader array of resistance genes is involved in deltamethrin resistance in Kor-R and F1 hybrids than those in TWB-R (Table 2).

**Table 2.** Bioassay results for susceptible, resistant, and hybrid strains of *Helicoverpa armigera*.

| Strains | Deltamethrin |  |  |  |
| --- | --- | --- | --- | --- |
|  | LD<sub>50</sub> (μg/larva) | χ<sup>2</sup> | RR<sup>s</sup> | RR<sup>r</sup> |
|  | (95% CI) | Log10 (Dose) |  |  |
| TWB-S | 0.0022 (0.0012–0.0037) | 40.4 | 1 | - |
| TWB-R | 0.0582 (0.0428–0.0743) | 16.6 | 26 | 1 |
| F1-hybrid (Kor-R (♂) × |  |  |  |  |
| TWB-R (♀)) | 15.4443 (7.8383–29.2836) | 38.9 | 7020 | 265 |
| Kor-R | 32.7093 (17.5580–59.2322) | 37.8 | 14,868 | 562 |

<sup>s</sup>Resistance ratio (RR<sup>s</sup>) = LD<sub>50</sub> of the resistant strain/LD<sub>50</sub> of the susceptible strain, TWB-S.763 <sup>r</sup>Resistance ratio (RR<sup>r</sup>) = LD<sub>50</sub> of the resistant strain/LD<sub>50</sub> of the low-level resistant strain, TWB-R.

At approximately the LD_50_ (0.05 µg) and LD_10_ values (0.01 µg) of deltamethrin against TWB-R, TWB-S mortality did not significantly differ between the PBO-treated and untreated groups; contrastingly, TWB-R mortality significantly differed. These consistent results were observed across various doses, indicating that the mortality rate increased when CYP was inhibited, thereby corroborating that CYP plays a major role in the development of resistance in TWB-R. Similar results were obtained for Kor-R, and the difference in mortality between PBO treatment and non-treatment was evident at 100 and 10 ppm, similar to the LC_50_ and LC_10_ values of deltamethrin. From these findings, it can be concluded that additional CYP (including CYP337B3) contributes to the development of insecticide resistance, although determining the exact extent of resistance in Kor-R remains challenging (Fig. 1).

**Fig. 1.**
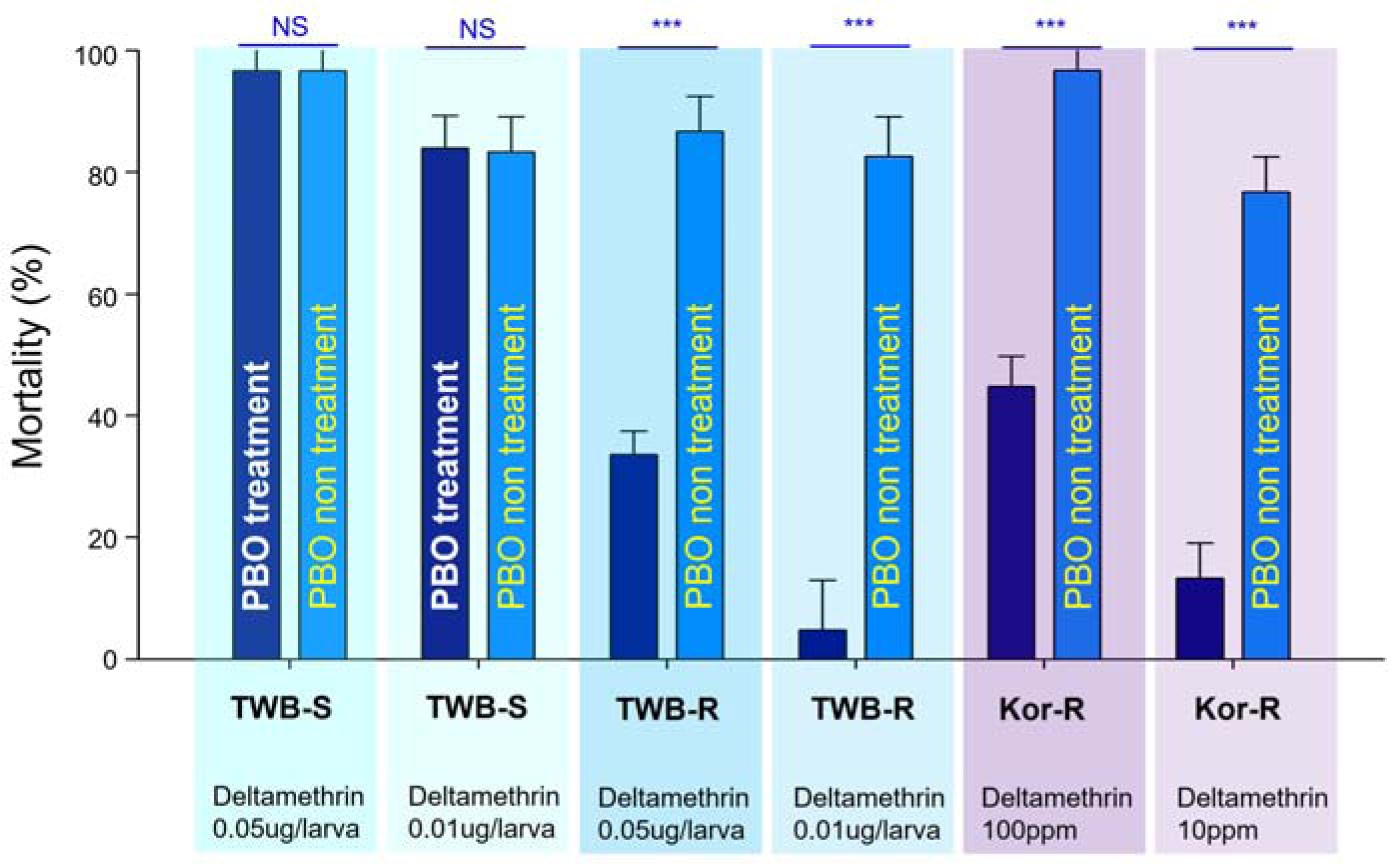
Synergist test results of a susceptible and two resistant *Helicoverpa armigera* strains against deltamethrin with or without 2 mM PBO as a cytochrome P450 inhibitor. Third-instar Australian susceptible (TWB-S) and resistant (TWB-R) *H. armigera* strain larvae were treated with two different doses (approximately LD_50_ and LD_10_) of deltamethrin with or without the synergist PBO. The Korean resistant strain (Kor-R) was also treated with two doses of deltamethrin. Mortality was assessed 2 days after treatment. PBO, piperonyl butoxide

### Mutation confirmation through the VGSC consensus genome of *H. armigera* and PCR

A graphic presentation linked to pyrethroid pesticide resistance is presented in Fig. 2a. The reference genome was aligned exclusively with 13 re-sequenced genomes at the VGSC gene loci. The total length of the VGSC gene sequence was 27,527 bp, comprising 34 exons and 33 introns (accession number MG674159.1). No site-specific mutations (V418, I947, L1023, D1556, and E1560) were observed throughout the complete sequence of VGSC in *H. armigera* [refer to the corresponding amino acid residues (ARK07244.1, length 2,042 amino acids) of *H. armigera*] (Fig. 2b). Since the consensus genome was sequenced on an individual basis, we reconfirmed the variants by PCR. At least three individuals per strain and population were analyzed to identify more detailed individual-level variations. Despite these efforts, no variants or mutations within VGSC were identified (Fig. 2c).

**Fig. 2.**
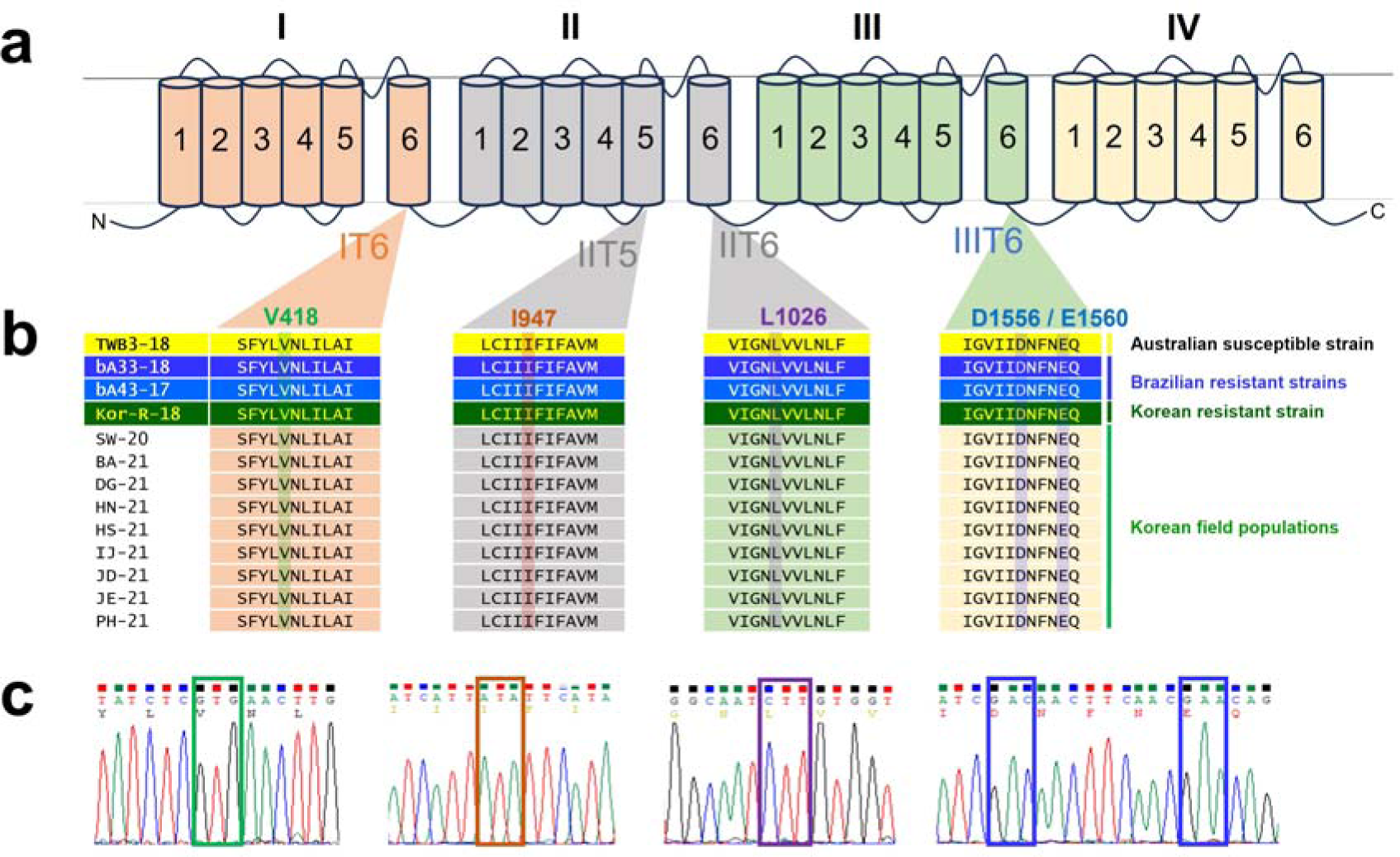
Schematic diagram of the five reported mutations based on pyrethroid insecticide resistance. **a** General pictorial view of four transmembrane domains (I–IV) and mutation positions (‘I–IV’ = domain number and ‘T 1–6’= transmembrane segment number). **b** Common mutation pattern of voltage-gated sodium channel of *Helicoverpa armigera*. The corresponding number of mutations reference to accession no. KY247121 (protein ID_ARK07244.1; total amino acids 2,042). Three mutations were reported (V421A/G, I951V, and L1029H; refer to GenBank accession no. GU574730) in the pyrethroid resistance mutation of *Helicoverpa zea* (Hopkins et al 2010). The two novel mutations, D1556 and E1560, were identified by Head et al. (1998). These two-point mutations were also functionally verified by Chen et al. (2017). **c** Representative chromatogram. The five reported mutations were reconfirmed by PCR.

### Whole-genome and re-sequencing analyses of H. armigera

To construct the primary reference genome sequence of *H. armigera,* the contig sequences of Kor-R were assembled using Nanopore long-read and Illumina short-read data (GCA_017165865.1). Subsequently, these contig sequences were refined to the chromosomal level using Hi-C data (GCA_026262555.1). The final genome sequence was 390.2 Mb in length with an N50 of 11.4 Mb and composed of 31 pseudo-chromosomes and 674 contig sequences. Repetitive DNA (23.83%; 92.91 Mb) corresponding to putative transposable elements was found in the *H. armigera* genome. The predicted number of genes in the *H. armigera* genome was 24,154. The percentages of GC content and GC skew at each chromosomal level are shown in Fig. 3a. Despite their distinct feeding habits, *H. armigera* (GCA_026262555.1) and *H. zea* (GCA_002150865.1) genomes shared high levels of orthology, indicating a common evolutionary origin (Fig. 3b). As this study focused on insecticide resistance, further details on comparative genome research are not provided.

**Fig. 3.**
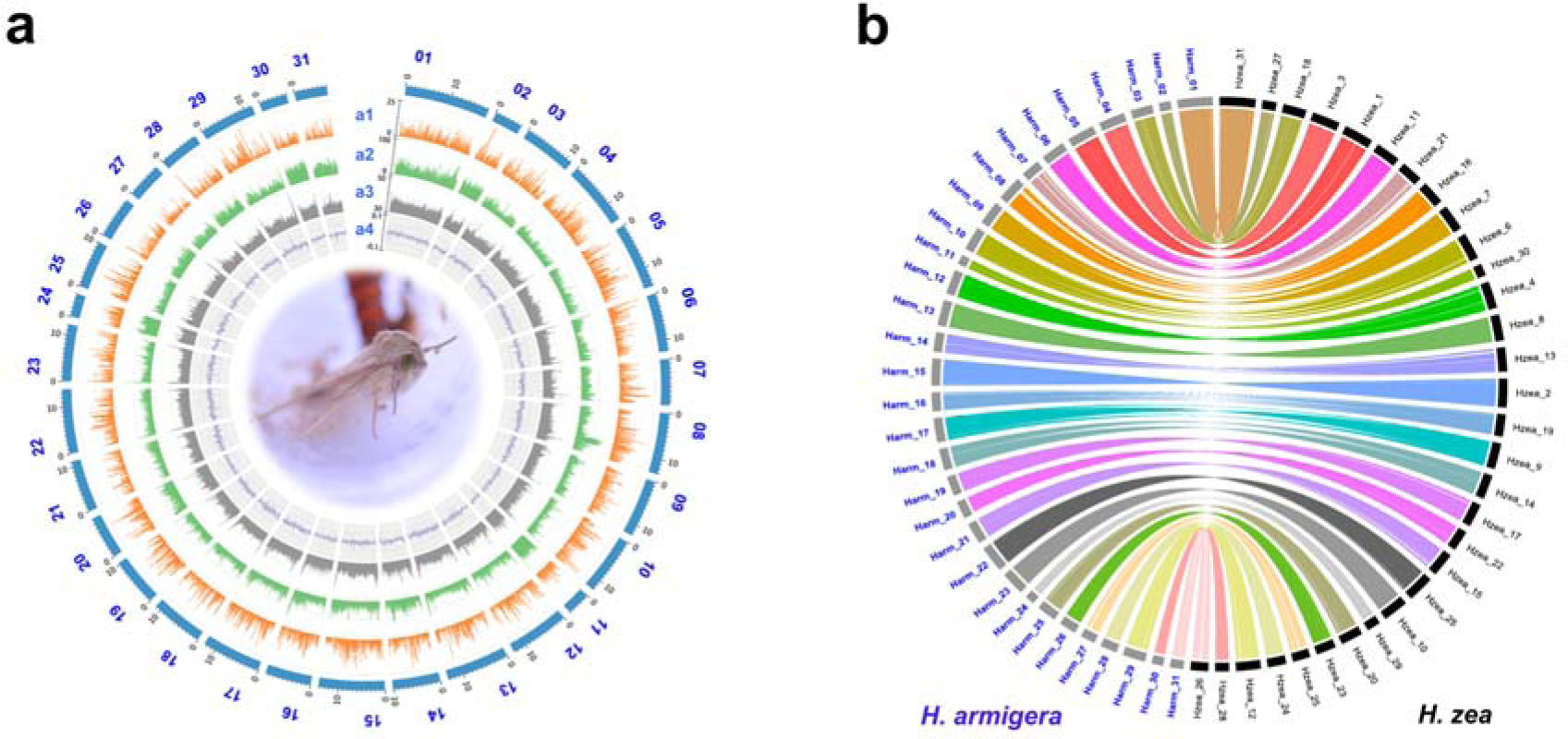
Integration of genomic and expression data. **a** distribution of genes (number of genes per 100 Kb) in each chromosome. (a2) % of repeat distribution. (a3) % GC content. (a4) GC skew [(G - C)/(G + C)]. **b** Genome synteny analysis of *Helicoverpa armigera* and *Helicoverpa zea*. The considered window size was 100 Kb. Synteny analysis was conducted using MCScanX and visualized using Circos v0.69-9.

The re-sequencing results of the genomes (*n=*13) revealed a total of 3,015,579 bp variants (SNPs and indels) in TWB-S and 3,308,578 bp in bA33 (Table 3). Notably, Kor-R exhibited slightly fewer variants than those of other resistant strains, including those collected from Brazilian, Australian, and Korean field populations (Table 3). Additionally, various categories of mutations, including intergenic, synonymous, and non-synonymous mutations, were identified (Tables S2).

**Table 3.** Re-sequencing data and mutation type categories in the susceptible, resistant, and Korean field populations.

| Sample code | Sample sources | Total variants (SNPs+ Indels) | <sup>a</sup>SNPs | <sup>b</sup>Indels | Genic variants | Intergenic variants | Total length (bp) |
| --- | --- | --- | --- | --- | --- | --- | --- |
| TWB-S | Susceptible strain from Australia | 3,015,579 | 2,521,925 | 493,654 | 926,312 | 2,089,267 | 389,788,739 |
| bA33 | Resistant strain from Brazil | 3,308,578 | 2,762,822 | 545,756 | 1,017,018 | 2,291,560 | 389,759,414 |
| bA43 | Resistant strain from Brazil | 3,098,092 | 2,590,939 | 507,153 | 952,532 | 2,145,560 | 389,782,873 |
| Kor-R | Resistant strain from Korea | 2,595,043 | 2,168,783 | 426,260 | 800,098 | 1,794,945 | 389,867,883 |
| BA-21 | Buan, Korea | 3,566,291 | 2,976,583 | 589,708 | 1,090,032 | 2,476,259 | 389,733,990 |
| DG-21 | Daegwallyeong, Korea | 3,570,286 | 2,982,026 | 588,260 | 1,092,966 | 2,477,320 | 389,705,348 |
| HN-21 | Haenam, Korea | 3,566,817 | 2,976,169 | 590,648 | 1,090,825 | 2,475,992 | 389,733,451 |
| HS-21 | Hoengseong, Korea | 3,361,495 | 2,806,959 | 554,536 | 1,029,476 | 2,332,019 | 389,739,813 |
| IJ-21 | Injae, Korea | 3,567,847 | 2,979,529 | 588,318 | 1,091,643 | 2,476,204 | 389,708,178 |
| JD-21 | Jindo, Korea | 3,604,152 | 3,007,730 | 596,422 | 1,103,022 | 2,501,130 | 389,714,678 |
| JE-21 | Jeongeup, Korea | 3,569,699 | 2,980,072 | 589,627 | 1,091,811 | 2,477,888 | 389,733,728 |
| PH-21 | Pohang, Korea | 3,384,286 | 2,834,043 | 550,243 | 1,037,639 | 2,346,647 | 389,733,420 |
| SW-20 | Suwon, Korea | 3,599,720 | 3,002,322 | 597,398 | 1,101,587 | 2,498,133 | 389,708,844 |

<sup>a</sup>single nucleotide polymorphisms; <sup>b</sup>insertions and deletions.

### Reference genome-based DEG analysis

A total of 45 RNA-seq datasets were obtained from three different tissue samples (fat body, gut, and the rest of the body) of one susceptible and three resistant *H. armigera* strains (Fig. 4). DEG analysis was carried out between insecticide-resistant strains (TWB-R, Kor-R, Kor-T, and bA43) and a susceptible strain (TWB-S) in tissues, based on the expression pattern using hierarchical clustering with Euclidean distance (Fig. 4a). The clustering revealed patterns of up-regulated and down-regulated transcripts, providing insights into the differences in gene expression between susceptible and resistant strains. While the differences between resistance and expression levels were not correlated, Kor-R and bA43, which showed moderate and high levels of resistance, respectively, exhibited differential expression of various genes compared to TWB-S, a susceptible strain (Fig. 4b).

**Fig. 4.**
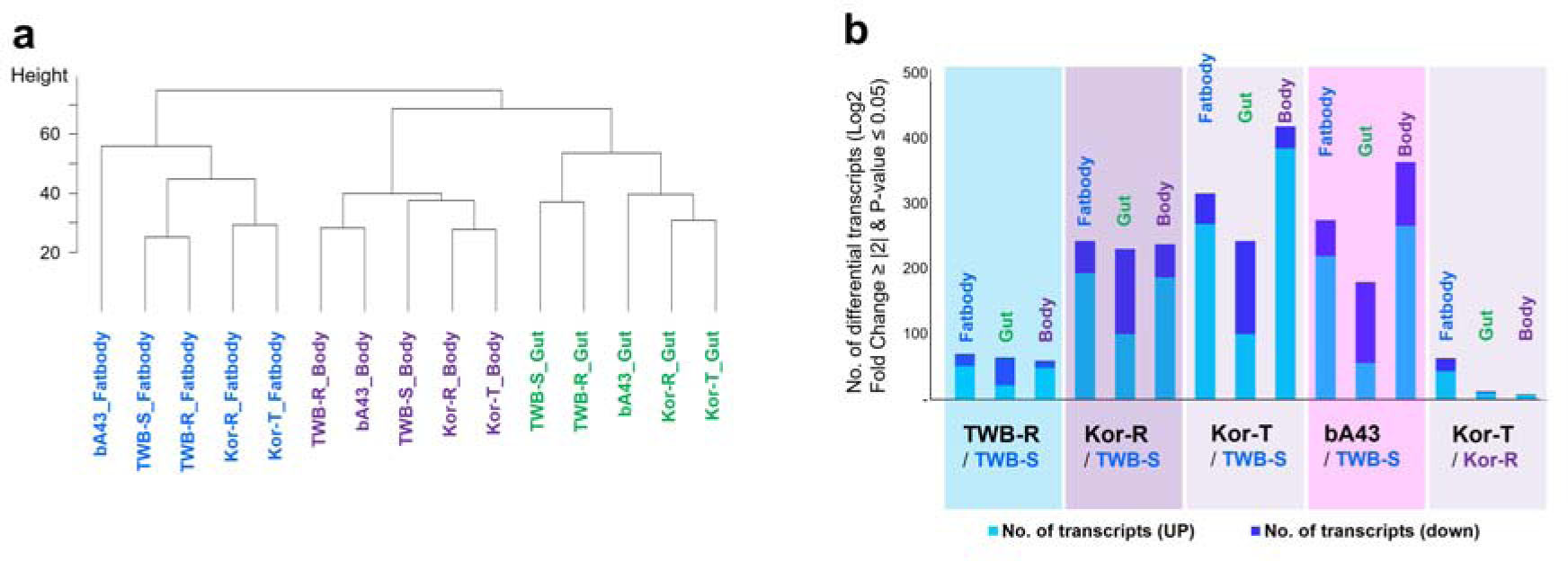
Number of differentially expressed coding RNAs (up- and downregulated) based on the strains or tissues. **a** Hierarchical clustering of significant Euclidian distances in *Helicoverpa armigera*. The cluster represents differential transcripts compared between susceptible (TWB-S) and insecticide-resistant strains (TWB-R, Kor-R, Kor-T, and bA43). **b** Comparisons of differentially expressed transcripts in insecticide-resistant *Helicoverpa armigera* strains (TWB-R, Kor-R, Kor-T, and bA43) compared with those of the susceptible strain (TWB-S). Black color indicates fat body; blue color indicates the rest of the body; green color indicates intestinal tissue.

Comparing the overall transcript expression patterns, the differences between tissues were more distinct than those between strains (TWB-R, TWB-S, Kor-R, Kor-T, and bA43) with different resistance levels. Notably, the gene expression pattern in the rest of the body was more similar to that in the gut than that in the fat body (Fig. 5). A comparison of strains with different insecticide resistance levels revealed that the numbers of significantly expressed genes (*p* ≤ 0.05; up- and downregulated genes) relative to the susceptible strain were 193, 712, and 819 for TWB-R, Kor-R, and bA43, respectively (Table S5). This suggests that gene expression levels vary with resistance (low to high resistance). Differences in gene expression considering susceptibility (TWB-S) varied depending on the resistance level.

**Fig. 5.**
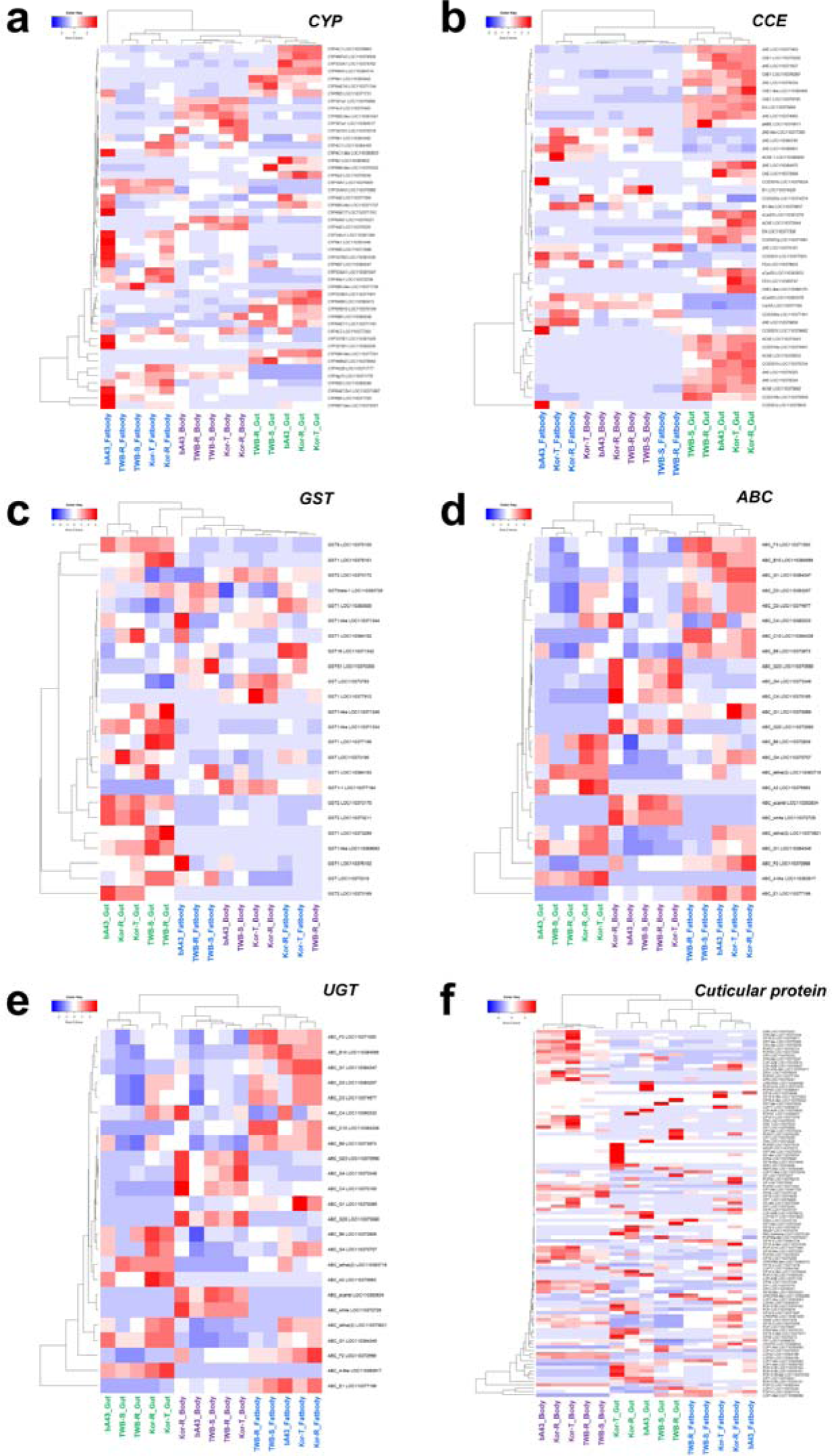
Expression pattern of major coding genes, including **a** cytochrome P450 (CYPs), **b** carboxylesterases (CCEs), **c** glutathione S-transferase (GST), **d** ABC transporter (ABC), **e** UDP-glycosyltransferase (UGT), and **f** cuticular protein (CP). The heat map shows differentially expressed genes between a susceptible and three resistant strains of *Helicoverpa armigera*.

Because gene expression differed depending on the level of insecticide resistance, detailed expression differences were compared, focusing on six major detoxification gene families, including CYP, CCE, GST, ABC transporter, and UGT, as well as cuticular protein genes (Fig. 5). Some detoxification genes, such as CYP333B3 |LOC110377401|, were highly expressed in gut tissue compared with other tissues (fat body and the rest of the body). In contrast, CYP337B1 (LOC110381029) and CYP321B1 (LOC110382836) were expressed at higher levels in fat body than in other tissues (Fig. 5a). Notably, cytochrome 6 B family members (CYP6B5– 7) exhibited varied expression patterns, sometimes being expressed in both the gut and fat body tissues (Fig. 5). Similarly, certain metabolic enzymes were expressed differentially in all three tissue categories. These genes have tissue-specific expression patterns, with different members of their families expressed more prominently in particular tissues, such as the gut, fat body, or both (Fig. 5b-f). These expression patterns may have functional implications for the roles of these genes in different physiological processes within cells.

For CCE, one gene (CCE001j, LOC110379835) was overexpressed in bA43. However, for other detoxification enzymes, the types of genes expressed in each tissue differed without showing a significant difference in expression depending on the tissue. The expression level of cuticular protein genes was particularly high in bodies containing exoskeletal tissue, and its expression appeared to increase in Kor-T and Kor-R following treatment with a sublethal deltamethrin dose. The comparison of Kor-R and Kor-T expression before and after deltamethrin treatment is summarized separately in Fig. 7.

**Fig. 6.**
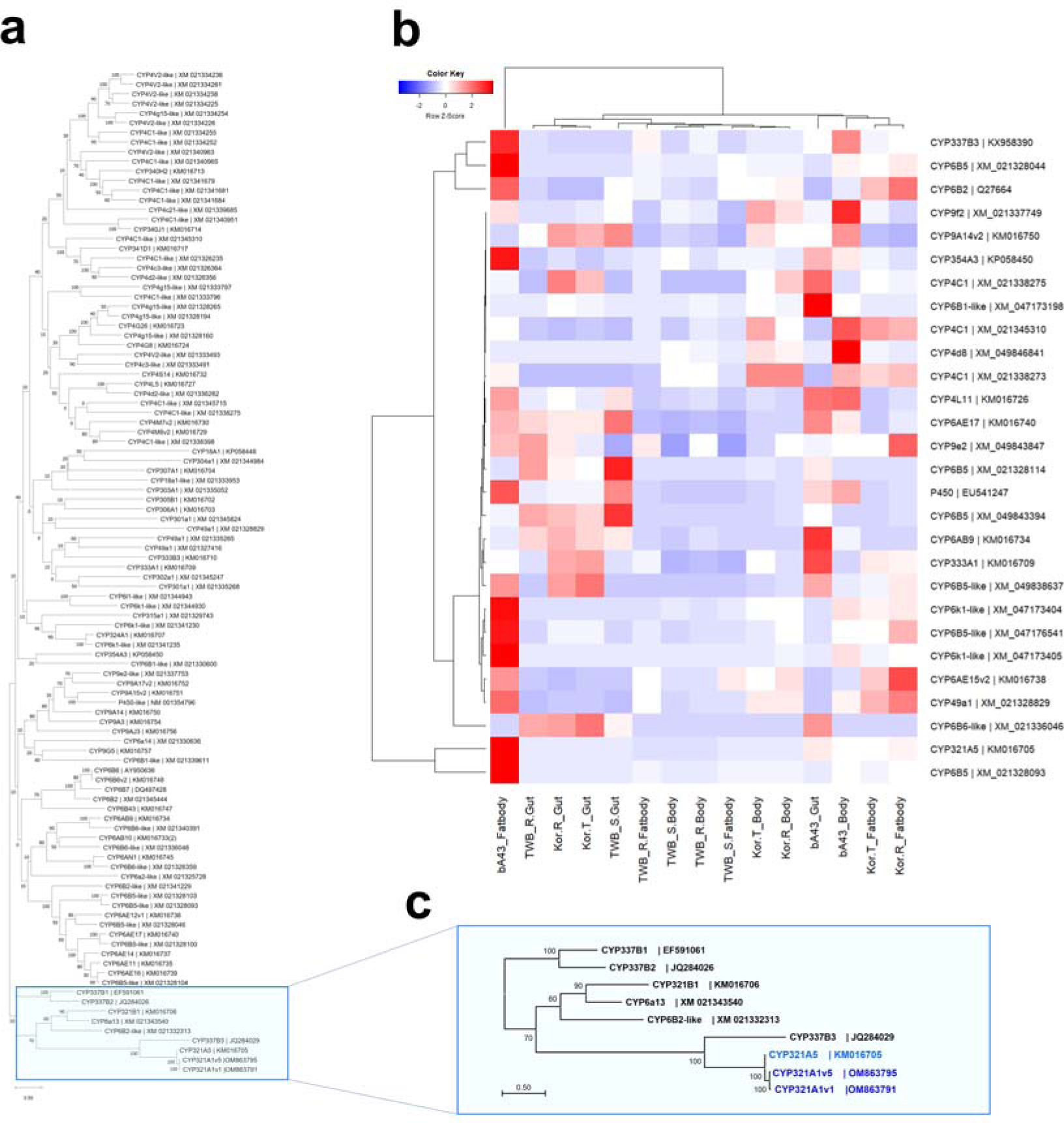
Phylogeny and expression pattern of major cytochrome P450 detoxification genes (CYPs) based on UniGene. The heat map shows differentially expressed genes between a susceptible and three resistant strains of *Helicoverpa armigera*. CYP genes with mixed expression levels showed higher expression in resistant strains than that in the susceptible strain. Color represents high (deep red) and low (light blue) CYP gene expression. Sample codes and gene symbols are shown at the bottom and on the right side of the heatmap, respectively. Sample codes: bA43, high pyrethroid resistance Brazilian strain; Kor-R, resistant Korean strain without treatment; Kor-T, resistant Korean strain with deltamethrin treatment; TWB-R, resistant Australian strain; TWB-S, susceptible Australian strain. The extraction order for samples was gut first, followed by fat body and the rest of the body.

**Fig. 7.**
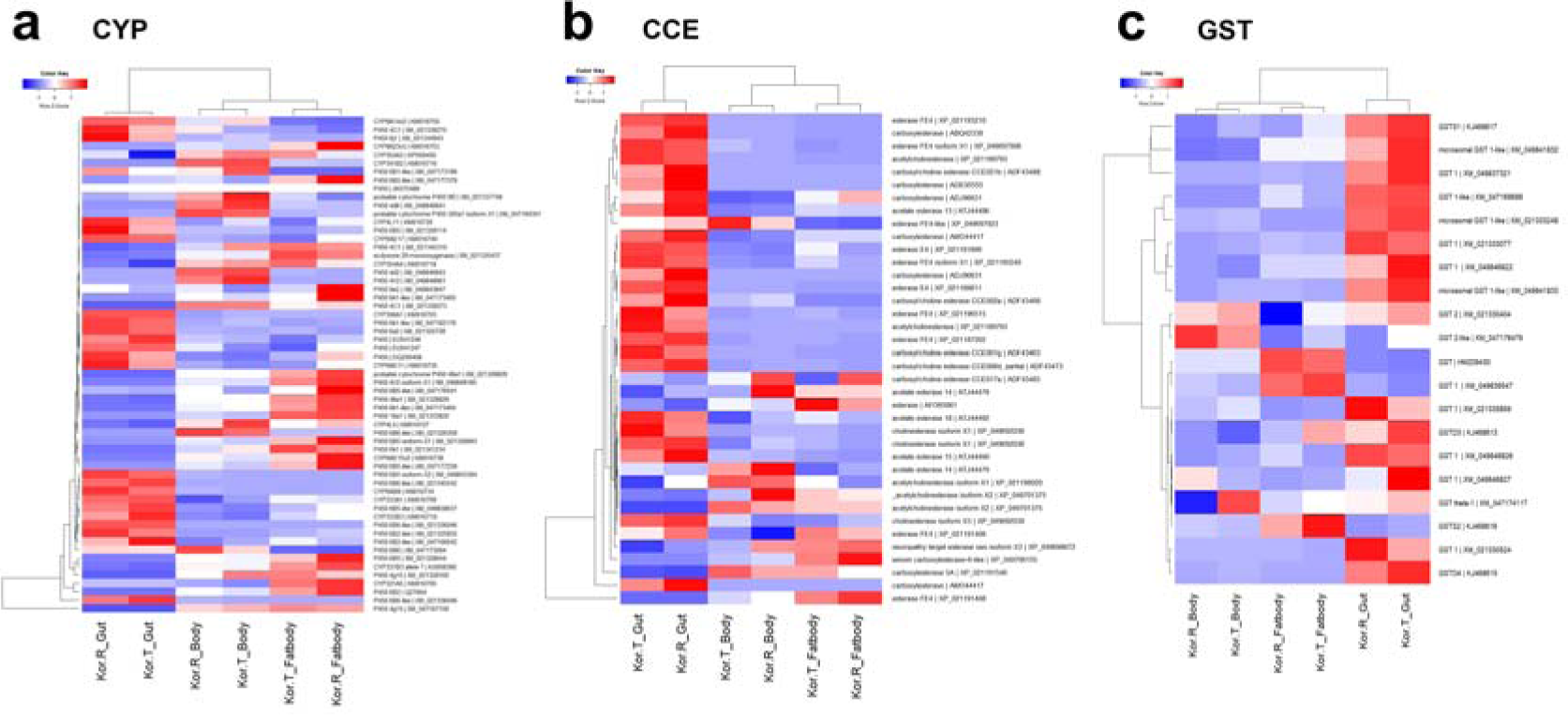
Comparison of expression patterns for three major detoxification gene families in artificially induced (Kor-T-treated) and normal Kor-T (Kor-R-untreated) strains, including **a** cytochrome P450 (CYP), **b** carboxylesterases (CCE), and **c** glutathione S-transferase (GST) based on the UniGene set. The heat map shows differentially expressed genes between treated (Kor-T) and non-treated resistant *Helicoverpa armigera* (Kor-R) strains

### UniGene set-based DEG analysis

The expression patterns of major detoxification genes, specifically P450s, in *H. armigera,* based on UniGene, form a cluster with CYP337B, including the respective CYP3 subfamily (Fig. 6a and c). Furthermore, considering the high similarity (ca. 96–97%) of the CYP6B2-like (XM021332313) gene with the CYP321A genes (Fig. 6c) and the absence of CYP337B3, which is involved in pyrethroid insecticide resistance, from the reference genome database (Harm1.0), a UniGene set was created (detailed in the materials and methods section) and the DEGs were reanalyzed using the UniGene sets. The color coded a heat map visually represents the expression levels of differentially expressed genes between susceptible and resistant strains. The color code on the heat map demonstrates expression levels, with deep red indicating higher expression and light blue indicating lower expression of P450 genes (Fig. 6b). The presence of CYP321A1 was confirmed by comparing the genomic information of the TWB3 strain (GCA_002156985.1), which is considered the parent of TWB-S and TWB-R, with that of the Kor-R strain (GCA_026262555.1) obtained in this study, confirming its presence only in the Kor-R genome. This gene, located on chromosome 6, has an open reading frame of approximately 1.5 kb, no introns, and two identical genes in a row with a gap of approximately 13 kb (GCA_026262555.1). Variants were also present, with two identified in Kor-R: CYP321A1v1 and CYP321A1v2 (OM863791.1 and OM863792.1, respectively), while CYP321A1v3 and CYP321A1v4 were detected in bA43 (OM863793.1 and OM863794.1, respectively). Similarly, a CYP321A1v5 variant was detected in another Brazilian resistant strain, bA33 (OM863795.1). The presence of CYP321A5, along with five members of CYP321A1v1–v5, was confirmed using gene-specific primers (Table S1). As a result, CYP321A5 commonly exist in all tested samples, but CYP321A1 specifically detected via conventional PCR and LAMP in other resistant and Korean field populations (Figure S1). The exact function of this gene and the characteristics of each variant remains unknown, and research is ongoing.

### Comparative study of artificially induced (Kor-T-treated) and normal Kor-T (Kor-R-untreated) strains

This study identified a significant number of DEGs by comparing an artificially induced Kor-R strain (Kor-T-treated) to an untreated resistant strain (Kor-R-untreated). These DEGs were further categorized and analyzed based on their up- or downregulation status. When we compared the Kor-T and Kor-R strains, the highest number of DEGs (*n* = 63) was observed in fat body, among which 43 and 20 were up- and downregulated, respectively.

In the comparison of the Kor-T and Kor-R strains, differentially expressed genes were identified in the CYP, CCE, and GST gene families. Among the 59 highly expressed CYP genes, CYP337B3 (KX958390), CYP321A5 (KM016705), and CYP6B2 (Q27664) showed higher expression in fat body than in the rest of the body and gut (Fig. 7a). Concerning the CCE gene family, 38 genes were highly expressed in the gut tissue when Kor-T was compared with Kor-R. Notably, one specific gene, acetylcholine esterase (XMP_021189793), was highly expressed in the gut of Kor-T compared with that of Kor-R (Fig. 7b). Regarding the GST gene family, a substantial number of GST genes (32) were highly expressed when comparing Kor-T with Kor-R. In particular, the GST 1 isoform X1(XM049846822) showed higher expression in Kor-T (deep red color) than that in Kor-R in gut (less intense red color) (Fig. 7c). Overall, within the three gene families (CYP, CCE, and GST), the gut tissue exhibited the highest gene expression levels compared to the fat tissue and the rest of the body (Fig. 7a–c).

## DISCUSSION

Pyrethroid insecticides have been extensively used owing to their perceived safety and minimal side effects on other mammals (Palmquist et al., 2012; Mueller-Beilschmidt, 1990;Wang et al. 2018). Specific point mutations, such as V421M/A/G, I951V, and L1029H, have been identified in numerous lepidopteran species and are associated with pyrethroid resistance, including heliothine taxa such as *H. armigera*, *H. zea*, and *Chloridea virescens* (Head et al. 1998; Hopkins and Pietrantonio 2010). In addition, geographic variability exists in the occurrence of these mutations. In some insect populations, these mutations may contribute to resistance, whereas in others, they may or may not occur. Resistance in all cases is not solely due to point mutations in VGSC genes, as some insect populations may exhibit resistance mediated by metabolic resistance genes. The presence of specific metabolic resistance genes, such as CYP337B3 in *H. armigera* samples, has been observed. A recent study by Ni et al. (2023) assessed 1,439 individuals but did not find point mutations associated with resistance in the examined individuals. Instead, they observed the presence of metabolic resistance genes, specifically CYP337B3, in *H. armigera* samples (Ni et al. 2023).

The molecular mechanisms underlying insecticide resistance are complex, encompassing mutations, SNPs, indels, and variations in expression levels (Nauen et al. 2022). The present study involved the whole-genome *de novo* assembly of Kor-R (GCA_026262555.1), the re-sequencing of 13 genomes from both susceptible and resistant individuals from field populations, and the identification of SNPs and indels in their genomes. Although a significant number of SNPs and indels were detected, none of these specific variants strongly correlated with insecticide resistance in *H. armigera*. This suggests that the genetic basis of insecticide resistance in this population may be more complex and may involve multiple genes or regulatory elements rather than specific individual mutations.

High-throughput RNA-seq was performed to further elucidate the molecular mechanisms underlying insecticide resistance. The RNA-seq data showed that TWB-R, characterized by a low degree of resistance, exhibited the lowest differential expression compared to TWB-S. TWB-R strain was selected based on fenvalerate resistance (∼40 times) within the same population of susceptible TWB-S strain. The primary cause of TWB-R resistance is the detoxifying enzyme, encoded by CYP337B3, which directly degrades fenvalerate (Joußen et al. 2012; Joußen and Heckel 2021). Resistant strains, such as Kor-R and bA43, with high levels of resistance, also harbor the detoxifying enzyme, encoded by the same gene, CYP337B3 (Durigan et al. 2017; Kim et al. 2018). In the same context, Kor-R, Kor-T, and bA43 showed significantly different expression patterns from TWB-S, unlike TWB-R. Additionally, Kor-R is not only genetically distant but also has a higher level of resistance due to factors other than CYP337B3 (Durigan et al. 2017; Kim et al. 2018), consistent with the previous findings in Chinese populations (Ni et al. 2023). Bioassay results showed a clear difference in the resistance ratio (RR) between Kor-R and TWB-S, as well as TWB-R. The F1 hybrid strain (Kor-R male and TWB-R female) was considered, because females can transmit genetic material to the next generation. The degree of resistance of the F1 hybrid was similar to that of susceptibility, indicating that half the level of resistance observed in Kor-R can be attributed to the transmission of genes related to metabolism to the next generation.

Transcriptional regulation by trans- and/or cis-factors, as well as copy number variation, influences the overexpression of specific CYPs (Zanger and Schwab 2013; Bu et al. 2015; Balabanidou et al. 2018). In the present study, variations in the copy number of coding RNA were observed in the susceptible and resistant strains of *H. armigera*. Several studies have documented CYP overexpression in pyrethroid-resistant strains of *H. armigera* (Brun-Barale et al. 2010; Trapnell et al. 2013). The CYPs (CYP9A12, CYP9A14, and CYP6B7) and insecticide-degradative CYP3 clans are divided into CYP6 and CYP9 (Feyereisen 2012), which are constitutively overexpressed in the fat body of fenvalerate-resistant strains compared to susceptible strains (Wu et al. 2011; Xu et al. 2016). Considerable research has shown that CYP3 is involved in insecticide metabolism via direct detoxification processes (Nauen et al. 2022). Furthermore, the CYP332A locus on chromosome 15 and its duplications have been identified in *H. armigera.* Notably, between the two duplicated copies (CYP332A1 and CYP332A2), the transposable element may provide insights into the process involved in duplication (Sezutsu et al. 2013). CYP332A1 is also significantly expressed in all six field strains surveyed in China (Joußen and Heckel 2021). However, Xu et al. 2016 observed no positive relationship between the resistance and expression levels of CYP genes (Xu et al. 2016).

Several CYP genes were investigated in the current study, including the CYP3 subfamily genes CYP337B1, CYP337B2, and CYP337B3, as well as the identified variant CYP321A1v5, which may be associated with deltamethrin resistance. In a recent study, Ni et al. (2023) suggested that point mutations in the VGSC gene are not major contributing factors to current pyrethroid resistance in *H. armigera* populations (Ni et al. 2023). CYP337B3 has been observed at significantly higher frequencies than CYP337B1 and CYP337B2 in most *H. armigera* populations. The higher frequency of CYP337B3 suggests that possessing this gene could provide a selective advantage, likely owing to its involvement in pyrethroid resistance. This implies that individuals with CYP337B3 may have better survival or reproduction rates when exposed to pyrethroid insecticides (Ni et al. 2023). The current study also suggests that the mutation is not only a factor of resistance but also a metabolic gene, specifically CYP337B3, with additional CYP321A1v1–v5 variants being the main candidates.

However, in certain field-derived resistant *H. armigera* populations in China, fenvalerate resistance was unrelated to CYP337B3 (Han et al. 2015), suggesting that CYP-mediated resistance in *H. armigera* is a complex phenomenon involving multiple contributing factors. Previous studies have also indicated that CYP, such as chimeric CYP (CYP337B3), is associated with *H. armigera* insecticide resistance (Scott and Wen 2001; Joußen and Heckel 2021). The presence of CYP337B3, resulting from unequal crossover between CYP337B2 and CYP337B1, has been identified in *H. armigera* populations resistant to fenvalerate in Australia (Joußen et al. 2012), cypermethrin in Pakistan (Rasool et al. 2014), deltamethrin in Korea (Kim et al. 2018), several field populations in Brazil (Durigan et al. 2017), and worldwide (Joußen and Heckel 2021).

Complex CYP-mediated insecticide resistance mechanisms involve enhanced insecticide metabolism, resulting in fewer toxic byproducts. The overexpression of particular CYP genes can accelerate the detoxification of insecticides and impart resistance. For example, increased CYP6BQ23 expression is associated with higher levels of deltamethrin detoxification and resistance in populations of the pollen beetle (*Meligethes aeneus*) (Nauen et al. 2022). Various P450 enzymes can metabolize a number of insecticides, including CYP6M2, which metabolizes pyrethroids and DDT (Mitchell et al. 2012), whereas some P450 enzymes play specialized roles in the metabolism of specific insecticides. Additionally, some P450s have functional redundancy, whereby multiple P450 enzymes within a subfamily can efficiently metabolize the same insecticide. CYP-mediated insecticide resistance is a complex and multifaceted phenomenon that involves various P450 enzymes with diverse substrate specificities.

Previous studies have demonstrated the *in vitro* activity of approximately 15 P450 enzymes from four different families (CYP6, CYP9, CYP321, and CYP337) against pyrethroids in various insect species (Feyereisen 2012; Joußen et al. 2012; Oakeshott et al. 2013). Subsequent investigations conducted in Pakistan revealed the presence of a novel chimeric P450 enzyme, identified as CYP337B3, which exhibited sequencing differences at the transcript variant level compared with those in an Australian study. Despite the geographical distance between Australia and Pakistan, both populations display insecticide resistance via the same mechanism involving CYP337B3. Consequently, the findings of Rasool et al. (Rasool et al. 2014) supported the notion that CYP337B3 is associated with insecticide resistance and pesticide degradation. Collectively, these studies contribute to understanding how certain enzymes play crucial roles in pesticide resistance across different geographical locations. However, it is important to note that the relationship between P450-mediated fenvalerate resistance and the CYP337B3 genotype have not been observed in Chinese *H. armigera* populations (Han et al. 2015). In Korean field populations, we identified CYP321, which may be involved in high resistance, and CYP337B3, which is involved in moderate resistance. In *P. xylostella*, the upregulation of CYP321E1 and CYP6BG1 has been observed in strains resistant to permethrin and chlorantraniliprole, respectively (Hu et al. 2014). Additionally, CYP321A8, CYP321A9, and CYP321B1 are responsible for *Spodoptera frugiperda* resistance to chlorantraniliprole (Bai-Zhong et al. 2020). The CYP337B3 enzyme of *H. armigera* has been shown to confer resistance to cypermethrin in Pakistan (Rasool et al. 2014) and fenvalerate in Australia (Joußen et al. 2012), which may be followed by a resistant strain of *H. armigera*. Several studies have shown that P450 confers insecticide resistance to susceptible insects (less expressed) than to resistant insects (overexpressed) (Yang et al. 2006; Fang et al. 2010). Overexpression can occur because of cis- or trans-acting transposable element insertions into the promoter region (Cariño et al. 1994). In addition, CYP duplication (Cariño et al. 1994; Schmidt et al. 2010) may also be a factor in protein overexpression.

Metabolic resistance to pyrethroids in *H. armigera* likely evolved through a rapid and significant change caused by a single amino acid mutation (L114F of CYP337B3 evolved by unequal crossover), providing a larger selective advantage than gradual stepwise improvements of the parental enzyme (Joußen and Heckel 2021). Metabolic factors synergize with PBO, which inhibits microsomal oxidases, a group of enzymes responsible for detoxifying foreign compounds in insects (Moores et al. 2009). The synergy experiment conducted in the present study demonstrated that the high levels of deltamethrin resistance observed in the resistant *H. armigera* strains could be significantly inhibited by PBO, with synergistic ratios ranging from 26-fold (TWB-R) to 14,868-fold (Kor-R). PBO is widely recognized as an inhibitor of P450 enzymes. These results further validate the crucial role of P450 enzymes in deltamethrin resistance in *H. armigera*. The fact that PBO effectively reversed resistance suggests that P450 enzymes are involved in the detoxification and metabolism of pyrethroid fenvalerate insecticides in *H. armigera* populations (Gunning et al. 1999). This finding is consistent with the results of the bioassay using PBO-based synergists, further supporting the role of detoxification enzymes in resistance. Another study investigated the effect of PBO on pyrethroid resistance-associated esterases in *H. armigera*, demonstrating the significance of PBO as a synergist in overcoming resistance mechanisms (Young et al. 2005).

The resistance mechanism in *H. armigera* depends on multiple factors responsible for its complexity. Recent genome projects have revealed an increasing number of paralogs (duplicated genes within a species) and orthologs (genes in different species with a common ancestral gene) in these subfamilies. However, their functionality remains poorly understood. The lepidopteran-specific CYP6AE subfamily has been previously implicated in pesticide resistance in *H. armigera*, and we observed the expression of the CYP6 clan. Notably, pesticide- and xenobiotic-metabolizing CYPs have been predominantly found in the CYP6 and CYP9 families of the CYP3 clan (Nauen et al. 2022). Incidentally, CYP9A paralogs are capable of metabolizing multiple pyrethroids in *H. armigera* (Tian et al. 2021). A previous study focused on the complete complement of CYP6B (four genes) and CYP9A (seven genes) in *H. armigera* and observed the heterologous expression of these CYP enzymes (in different hosts) in Sf9 cells and compared their functional activities. *In vitro* assays demonstrated that all CYP6B and CYP9A enzymes efficiently metabolized esfenvalerate pyrethroids (Shi et al. 2021). To date, little is known about the effects of long non-coding RNAs (lncRNAs) on the regulation of CYP expression (Yahouédo et al. 2017). Both coding and non-coding RNAs are believed to play a role in conferring resistance to certain substances, including pyrethroid insecticides. The current study did not observe a positive correlation between the expression levels of highly expressed CYP genes and the three tested resistant strains in the three different tissues analyzed (fat body, gut, and the rest of the body). Nevertheless, lncRNAs can act as either activators or repressors in the regulation of gene expression by directly binding to transcription factors or playing a role in DNA methylation. Histone modifications regulate the expression of CYPs at the transcriptional level, whereas lncRNAs can influence CYP expression at both transcriptional and post-transcriptional levels (Yahouédo et al. 2017). Further functional studies of lncRNAs are ongoing.

The resistance genes identified in the CYP321 subfamily may be associated with the high resistance observed in the Korean and Brazilian strains of *H. armigera*. To better understand the functions of coding and noncoding RNAs and their contributions to resistance mechanisms in insects, further research should be conducted. However, the current study identified five members (CYP321A1v1–v5) of the CYP321 variants, showing a potential role in the high resistance observed in Kor-R, bA43, and bA33. Additionally, we explored the molecular mechanisms and regulatory functions of these coding RNAs with respect to insecticide resistance. A limitation of the present study is the absence of functional analysis related to the resistance mechanisms of specific metabolic genes or coding proteins. Consequently, although certain genes were identified, their specific roles and mechanisms were not fully investigated or characterized. Furthermore, although it is not highly differently expressed, the study did not cover in-depth details on detoxification enzymes other than P450. Subsequence studies will address other detoxification enzymes, such as CCE.

Here, we conducted a genetic analysis focusing on five previously identified pyrethroid-associated point mutations within partial VGSC genes in the *H. armigera* populations. Notably, no point mutations associated with pyrethroid resistance were observed in the VGSC of *H. armigera*. While VGSC mutations are typically linked to insect resistance, we did not find such mutations in this population. Whole-genome analysis and re-sequencing of the Korean field populations did not identify specific SNPs or indels related to resistance mechanisms, suggesting that resistance in *H. armigera* populations is not driven by common genetic changes. Based on RNA-seq analysis of 45 RNA-seq datasets, coding RNAs were identified in susceptible and resistant *H. armigera* using systematic screening criteria. The six identified common detoxification enzyme genes were associated with regulatory roles in metabolic resistance, including CYP and cuticular proteins. Specifically, the presence of CYP337B3, along with the five members of CYP321A1 (v1 to v5), possibly involved in pyrethroid resistance, was confirmed using gene-specific primers. CYP321 subfamily genes, potentially linked to the high resistance in Korean and Brazilian *H. armigera* strains, were identified. Further in-depth research should be conducted on the functional characterization of the CYP321 subfamily and the regulatory mechanisms of overexpressed CYP genes, as well as their relationship with pyrethroid resistance mechanisms involving coding and lncRNAs in field populations of *H. armigera*. This study contributes to understanding the role of resistance mechanisms and lays the foundation for future research on *H. armigera* resistance management strategy.

## STATEMENTS AND DECLARATIONS

### Ethics approval

Not applicable.

### Consent to participate

Not applicable.

### Consent for publication

Not applicable.

### Availability of data and material

The genome assembly of *Helicoverpa armigera* (Kor-R strain) was submitted to NCBI with the assembly number GCA_017165865.1. The five identified CYP321A1 variants (v1– v5) have been deposited in NCBI under the accession numbers OM863791.1–OM863795.1. Data used in this study can be requested from the corresponding author.

### Code availability

Not applicable.

### Authors’ contributions

Conceptualization and designed experiments, J.K.; experiments, J.K., C.H., and J.J.; software, J.K., C.H., and M.R.; writing-original draft preparation, M.R. and J.K.; writing-review and editing, M.R., S.L., M.K., C.O., and J. K.; formal analysis and validation, M.R., S.L., and J.K.; resources, M.K., C.O., and J.K.; supervision, J.K.; project administration, M.K. and J.K. All authors read and approved the manuscript.

---

**Fig. S1.**
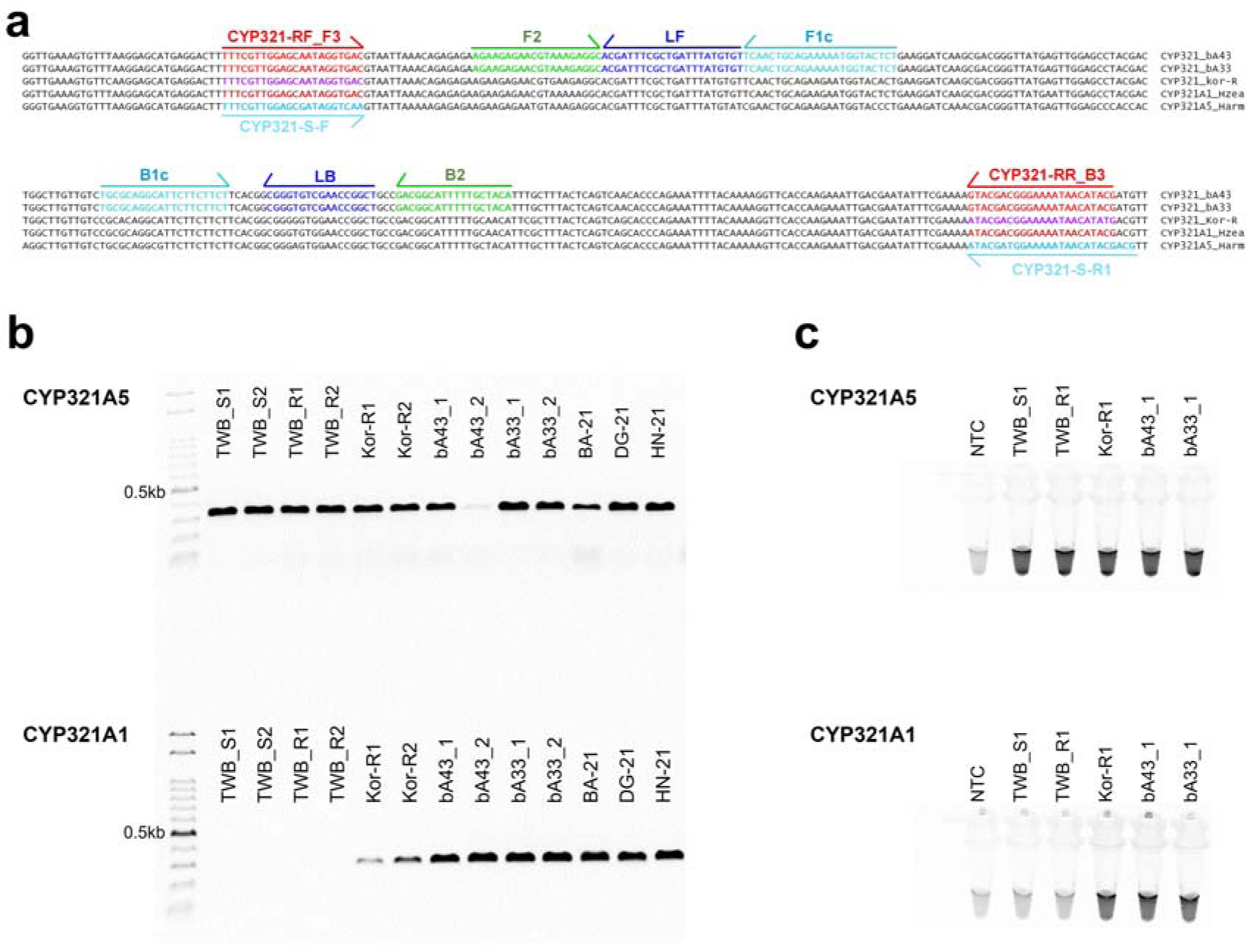
Development of Loop-mediated isothermal amplification (LAMP) and general PCR for specifically identifying the resistant CYP321A1 gene. **a**. LAMP-based six primers and primer-binding regions on the aligned partial gene sequences of CYP321A1 (from Korean resistant strain, Kor-R; two Brazilian resistant strains, bA43 and bA33; and *H. zea*) and CYP321A5. **b.** PCR results with each primer set using the same gDNA templates. **c**. LAMP assay result of insecticide resistant and susceptible strains, followed by visualization under ultraviolet light after treating with SYBR green. LAMP master mix and each set of primer were incubated with 50 ng of each gDNA at 61°C for 40 min. Symbols: TWB-S, Australian susceptible strain; TWB-R, Australian resistant strain; Kor-R, Korean resistant strain; bA43 and bA33, two Brazilian resistant strains; BA-21, field strain from Buan, Korea; DG 21, field strain from Daegwallyeong, Korea; HM 21, field strain from Haenam, Korea.

